# Targeting PFKFB3 alleviates cerebral ischemia-reperfusion injury in mice

**DOI:** 10.1101/518092

**Authors:** Olga Burmistrova, Ana Olias-Arjona, Tatiana Eremeeva, Dmitry Shishov, Kristina Zakurdaeva, Angeles Almeida, Peter O. Fedichev, Juan P. Bolaños

## Abstract

The glycolytic rate in neurons is low in order to allow glucose to be metabolized through the pentose-phosphate pathway (PPP), which regenerates NADPH to preserve the glutathione redox status and survival. This is controlled by 6-phosphofructo-2-kinase/fructose-2,6-bisphosphatase-3 (PFKFB3), the pro-glycolytic enzyme that forms fructose-2,6-bisphosphate, a powerful allosteric activator of 6-phosphofructo-1-kinase. In neurons, PFKFB3 protein is physiologically inactive due to its proteasomal degradation. However, upon an excitotoxic stimuli, PFKFB3 becomes stabilized to activate glycolysis, thus hampering PPP mediated protection of redox status leading to neurodegeneration. Here, we show that selective inhibition of PFKFB3 activity in neurons by the small molecule AZ67 prevents the NADPH oxidation, redox stress and apoptotic neuronal death caused by activation of glycolysis upon excitotoxic stimuli. Furthermore, *in vivo* administration of AZ67 to mice significantly alleviated the motor discoordination and brain infarct injury in the middle carotid artery occlusion ischemia/reperfusion model. These results show that pharmacological inhibition of PFKFB3 is a suitable neuroprotective therapeutic strategy for excitotoxic-related neurological diseases.

## INTRODUCTION

Glycolysis is widely considered a pro-survival metabolic pathway because it meets the energy needs of cells during mitochondrial bioenergetic stress [1]. However, in the brain tissue, different cell types show distinct metabolic preferences [2–4]. For instance, the metabolic use of glucose through glycolysis in neurons is normally very low, being mainly metabolized through the pentose-phosphate pathway (PPP), a metabolic route that contributes to the maintenance of neuronal redox status [5–8]. Astrocytes, in contrast, mainly obtain their cell energy needs from glycolysis, providing lactate as an oxidizable metabolic fuel to neurons [9], which obtain energy mainly by the oxidative phosphorylation [4].

A key factor that determines these metabolic features is 6-phosphofructo-2-kinase/fructose-2,6-bisphosphatase-3 (PFKFB3), a pro-glycolytic enzyme that is normally absent in neurons but abundant in astrocytes [7]. PFKFB3 activity produces fructose-2,6-bisphosphate (F2,6BP), a potent positive effector of the rate-limiting glycolytic enzyme, 6-phosphofructo-1-kinase (PFK1) [10, 11]. The absence of PFKFB3 protein in neurons is due to its continuous degradation after ubiquitylation by the E3 ubiquitin ligase anaphase-promoting complex/cyclosome-Cdh1 (APC/C-Cdh1) [7]. In fact, APC/C-Cdh1 activity is higher in neurons than in astrocytes [7]. Notably, under certain neuropathological conditions, such as during excitotoxicity, the activity of APC/C-Cdh1 in neurons is inhibited [12], which allows PFKFB3 protein stabilization in these cells [13]. Active neuronal PFKFB3 then stimulates glucose consumption through glycolysis, which results in a concomitant decreased PPP to cause redox stress and, eventually, apoptotic death [13].

Stroke is the leading neurologic cause of morbidity and mortality in developed countries [14]. While the molecular mechanisms underlying this complex pathological condition are not yet completely understood, a large body of experimental data suggest that excitotoxicity, leading to mitochondrial dysfunction and increased reactive oxygen species (ROS) are contributing factors [15–17]. Therefore, it appears reasonable that ameliorating the cascade of events triggered by excitotoxic stimuli might be a promising therapeutic strategy against stroke. Accordingly, we reasoned whether pharmacological inhibition of PFKFB3, by preventing the redox stress associated with glycolytic activation, would protect neurons from the apoptotic death upon excitotoxic insults. Here, we report that small molecule inhibitor of PFKFB3 in mouse primary cortical neurons is able to protect against the apoptotic death caused by N-methyl-D-aspartate (NMDA)- or glutamate-induced excitotoxic stimuli. Furthermore, we show that *in vivo* administration of this PFKFB3 inhibitor protects against motor discoordination and brain damage in a mouse model of brain ischemia/reperfusion.

## RESULTS

### *In vitro* characterization of two PFKFB3 inhibitors

First, we evaluated the efficacy of two known PFKFB3 inhibitors at inhibiting the ability of A549 cells to produce F2,6BP, namely AZ67 [18], and PFK158, an improved derivative of the widely used compound, 3-(3-pyridinyl)-1-(4-pyridinyl)-2-propen-1-one 1 (3PO) [19].

As shown in Figure 1A and 1B, both compounds (AZ67 and PFK158) were able to reduce the cellular levels of F2,6BP in a dose-dependent manner, with IC_50_ of 0.51 *µ*M and 5.90 *µ*M, respectively. Next, we investigated whether the reduction of cellular F2,6BP levels was a result of direct PFKFB3 inhibition. To do so, we used an enzymatic cell-free assay, which revealed that AZ67 inhibited the enzymatic activity of PFKFB3 with an IC_50_ of 0.018 *µ*M (Figure 1C), a value that is in accordance with previously published results [18]. However, surprisingly, PFK158 had no effect on PFKFB3 enzymatic activity at any of the concentrations tested (up to 100 *µ*M) (Figure 1D). Accordingly, although PFK158 is able to decrease F2,6BP (Figure 1B) and glycolytic flux [20], our data show that these effects are not due to PFKFB3 enzymatic inhibition. Since in this study we are focused specifically on PFKFB3 given its particular protein stability feature and potential impact on neurodegeneration, we did not consider PFK158 for further analyses.

**FIG. 1:**
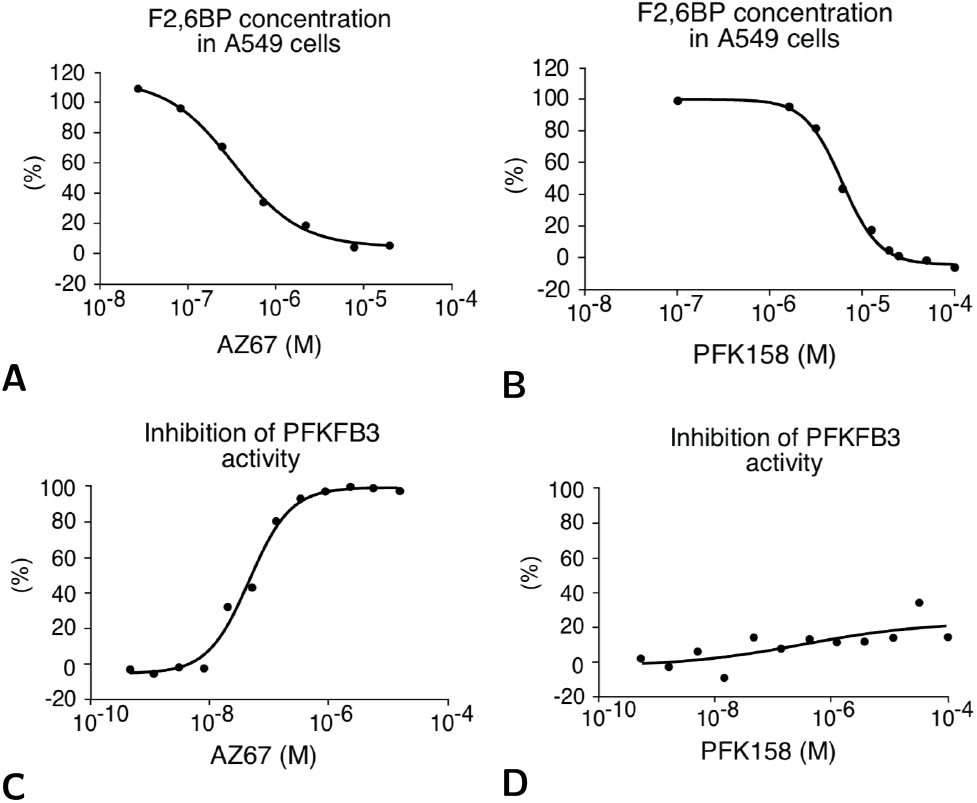
*In vitro* characterization of two PFKFB3 inhibitors. **A** Incubation of A549 cells with AZ67 (1 h) dose-dependently decreases F2,6BP concentration. **B** Incubation of A549 cells with PFK158 (1 h) dose-dependently decreases F2,6BP concentration. **C** Incubation of human recombinant PFKFB3 with AZ67 dose-dependently inhibits PFKFB3 kinase activity. **D** Incubation of human recombinant PFKFB3 with PFK158 does not inhibit PFKFB3 kinase activity.

### AZ67 protects neurons against proteasome inhibition and *β*-amyloid treatment

Since, in neurons, PFKFB3 is continuously degraded by the proteasome [7], we reasoned that the stabilization of PFKFB3 caused by proteasomal inhibition may trigger neuronal apoptosis.

As shown in Figure 2A, AZ67 lacks toxicity in the range 0.01-100 nM for 24 h in mouse cortical primary neurons. However, incubation of neurons with MG132, a widely used proteasomal inhibitor, significantly increased neuronal apoptosis (Figure 2A), an effect that was dose-dependently counteracted by AZ67 (minimum effective dose, 1 nM; maximum effect at 10 nM), suggesting the involvement of PFKFB3 activity in MG132-mediated neuronal death. To investigate if AZ67 protects neurons from the toxicity caused by PFKFB3 stabilization upon a different kind of stimulus, we next used the amyloidogenic fragment 25-35 of the amyloid-*β* peptide (*Aβ*_25−35_), known to activate glutamate receptors [21] and to inhibit Cdh1 [22], i.e. conditions that stabilize PFKFB3 [13]. Incubation of neurons with *Aβ*_25−35_ increased neuronal apoptosis (Figure 2A), and this effect was efficiently counteracted by AZ67 in a dose-dependent manner (minimum effective dose, 1 nM; maximum effect at 10 nM), thus suggesting that the excitotoxic effect of *Aβ*_25−35_ can, at least in part, be explained by PFKFB3 activation.

**FIG. 2:**
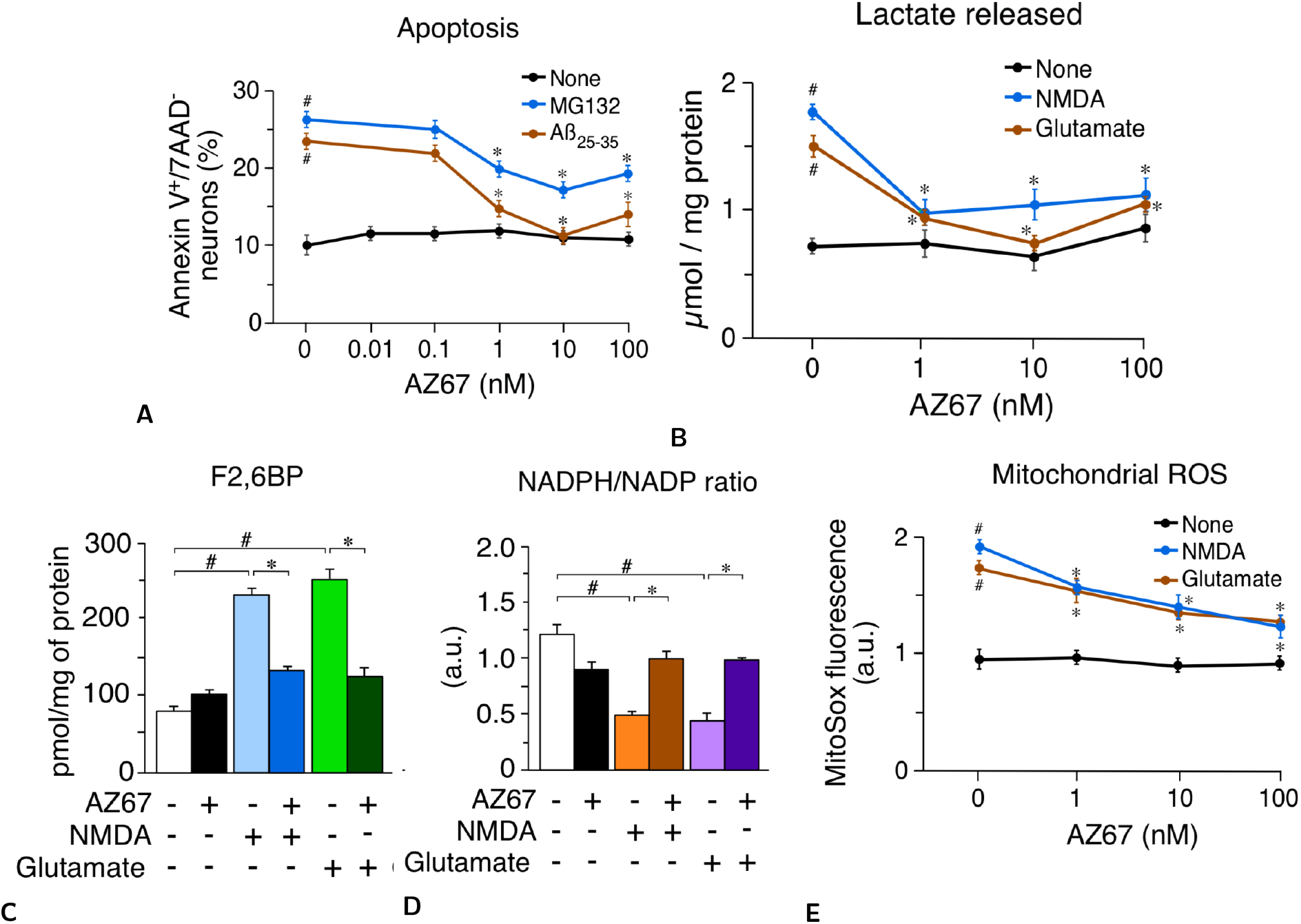
AZ67 protects neurons against proteasomal inhibition and *Aβ*_25−35_ treatment and rescues glycolytic activation and redox stress upon excitotoxic stimuli. **A** Incubation of mouse cortical primary neurons with AZ67 for 24 h revealed lack of toxicity. MG132 (10 *µ*M; 24 h) or *Aβ*_25−35_ (10 *µ*M; 24 h) increased neuronal apoptosis (compare MG132 or *Aβ*_25−35_ versus none values at 0 nM AZ67). Incubation of neurons with AZ67 together with MG132 or *Aβ*_25−35_, for 24 h, dose-dependently prevented apoptosis. **B** Incubation of neurons with AZ67 for 24 h revealed no effect on lactate release. Treatment of neurons with NMDA (100 *µ*M; 10 mins) or glutamate (100 *µ*M; 10 mins), followed by washout, increased lactate released after 24 h of incubation (compare NMDA or glutamate versus none values at 0 nM AZ67). Incubation of neurons with AZ67 for 24 h, after NMDA or glutamate was removed, dose-dependently prevented the increase in lactate release. **C** Incubation of neurons with AZ67 for 24 h revealed no effect on F2,6BP concentrations. Treatment of neurons with NMDA (100 *µ*M; 10 mins) or glutamate (100 *µ*M; 10 mins), followed by washout, increased F2,6BP after 24 h of incubation. Incubation of neurons with AZ67 (10 nM) for 24 h, after NMDA or glutamate was removed, prevented the increase in lactate release. **D** Incubation of neurons with AZ67 for 24 h revealed no significant effect on the NADPH/NADP ratio. Treatment of neurons with NMDA (100 *µ*M; 10 mins) or glutamate (100 *µ*M; 10 mins), followed by washout, decreased the NADPH/NADP ratio after 24 h of incubation. Incubation of neurons with AZ67 (10 nM) for 24 h, after NMDA or glutamate was removed, prevented the decreased NADPH/NADP ratio. **E** Incubation of neurons with AZ67 for 24 h revealed no effect on mitochondrial ROS. Treatment of neurons with NMDA (100 *µ*M; 10 mins) or glutamate (100 *µ*M; 10 mins), followed by washout, increased mitochondrial ROS after 24 h of incubation (compare NMDA or glutamate versus none values at 0 nM AZ67). Incubation of neurons with AZ67 for 24 h, after NMDA or glutamate was removed, dose-dependently prevented the increase in mitochondrial ROS. In all cases, data are mean ± S.E.M. values for n=3 independent culture preparations. #p<0.05 versus none at 0 nM AZ67; *p<0.05 versus the corresponding treatment at 0 nM AZ67 (ANOVA followed by the least significant difference multiple range test).

### AZ67 prevents glycolytic activation and redox stress upon excitotoxic stimuli in primary neurons

To test the ability of AZ67 to protect against the damage caused by an excitotoxic stimuli, neurons were subjected to a short-term incubation with NMDA or glutamate (100 *µ*M for 10 minutes) followed by a 24 h incubation in NMDA- or glutamate-free culture medium, a widely-used excitotoxic protocol [23]. In view that this treatment is known to trigger PFKFB3 protein stabilization [13], we assessed the glycolytic end-product, lactate, released to the culture medium after 24 h of incubation. As shown in Figure 2B, both NMDA- and glutamate-mediated excitotoxic stimuli were able to activate glycolysis, as judged by the increased lactate concentrations in the medium, an effect that was dose-dependently abrogated by incubation of neurons with AZ67 immediately after the excitotoxic stimuli during 24 h. The minimum concentration of AZ67 that showed to be fully efficient was 1 nM, although at 10 nM, AZ67 was maximally effective in the glutamate-mediated stimulus (Figure 2B). In good consistency with PFKFB3 protein stabilization [13], treatment of neurons with the excitotoxic stimuli increased the levels of the PFKFB3 product, F2,6BP, by almost ~ 3-fold (Figure 2C), indicating increased PFKFB3 enzymatic activity. Notably, the increased F2,6BP levels were fully abolished by AZ67 at 10 nM (Figure 2C). These data indicate that the pharmacological inhibition of PFKFB3 activity in neurons is sufficient to prevent excitotoxic stimuli-mediated activation of glycolysis. Since an increase in glycolysis leads to the impairment in the ability of neurons to regenerate NADPH through PPP activity [7], we next assessed the redox state of this cofactor. As shown in Figure 2D, stimulation of glutamate receptors triggered NADPH oxidation, a hallmark of PPP inhibition [7, 24], as judged by the decreased NADPH/NADP ratio, an effect that was abrogated by incubating neurons with AZ67 (10 nM). Given that PPP-mediated regeneration of oxidized NADPH is essential for preventing the redox stress in neurons [7, 13] that accompanies mitochondrial damage in several neurodegenerative diseases [25, 26], we next investigated mitochondrial reactive oxygen species (ROS). In good agreement with this notion, treatment of neurons with the excitotoxic stimuli promoted an increase in mitochondrial ROS (Figure 2E), and this effect was abolished by AZ67 (10 nM) (Figure 2E). Thus, inhibition of PFKFB3 activity upon an excitotoxic stimuli prevents the aberrant activation of glycolysis in neurons that leads to redox stress.

### The neuroprotective effect of AZ67 is lost by abrogating glycolytic activation

Next, we aimed to understand whether AZ67, by preventing the activation of glycolysis in neurons, could account for the neuronal death associated with the excitotoxic stimuli. To do so, neurons were incubated with glutamate of NMDA, as above, and apoptosis assessed by annexin V^+^/7AAD^−^ staining using flow cytometry.

As shown in Figure 3A, both types of excitotoxic stimuli significantly increased apoptotic neuronal death. Notably, this effect was dose-dependently prevented by AZ67, being 1 nM the minimum effective concentration and 10 nM the maximum dose showing protection (Figure 3A). A progressive loss of protection was observed at AZ67 concentrations ≥ 100 nM (Figure 3A). To address whether the neuronal protection exerted by AZ67 was a consequence of hampering the glycolytic activation, we assessed whether overexpression of the glycolytic enzyme, PFK1-muscle isoform (Figure 3B), was able to rescue AZ67-mediated neuroprotection. We focused on the muscle PFK1 isoform (PFK1-M) given its very low sensitivity to F2,6BP allosteric activation [27] and, hence, its independence on PFKFB3 levels to fully activate glycolysis [28]. Accordingly, neurons were first transfected with the full-length cDNA encoding for PFK1-M, and then subjected to the excitotoxic insults. As shown in Figure 3C, PFK1-M over-expression was able to abrogate the neuroprotection caused by AZ67 (10 nM). These results confirm that the neuroprotection exerted by AZ67 is a consequence of its ability to prevent glycolytic activation.

**FIG. 3:**
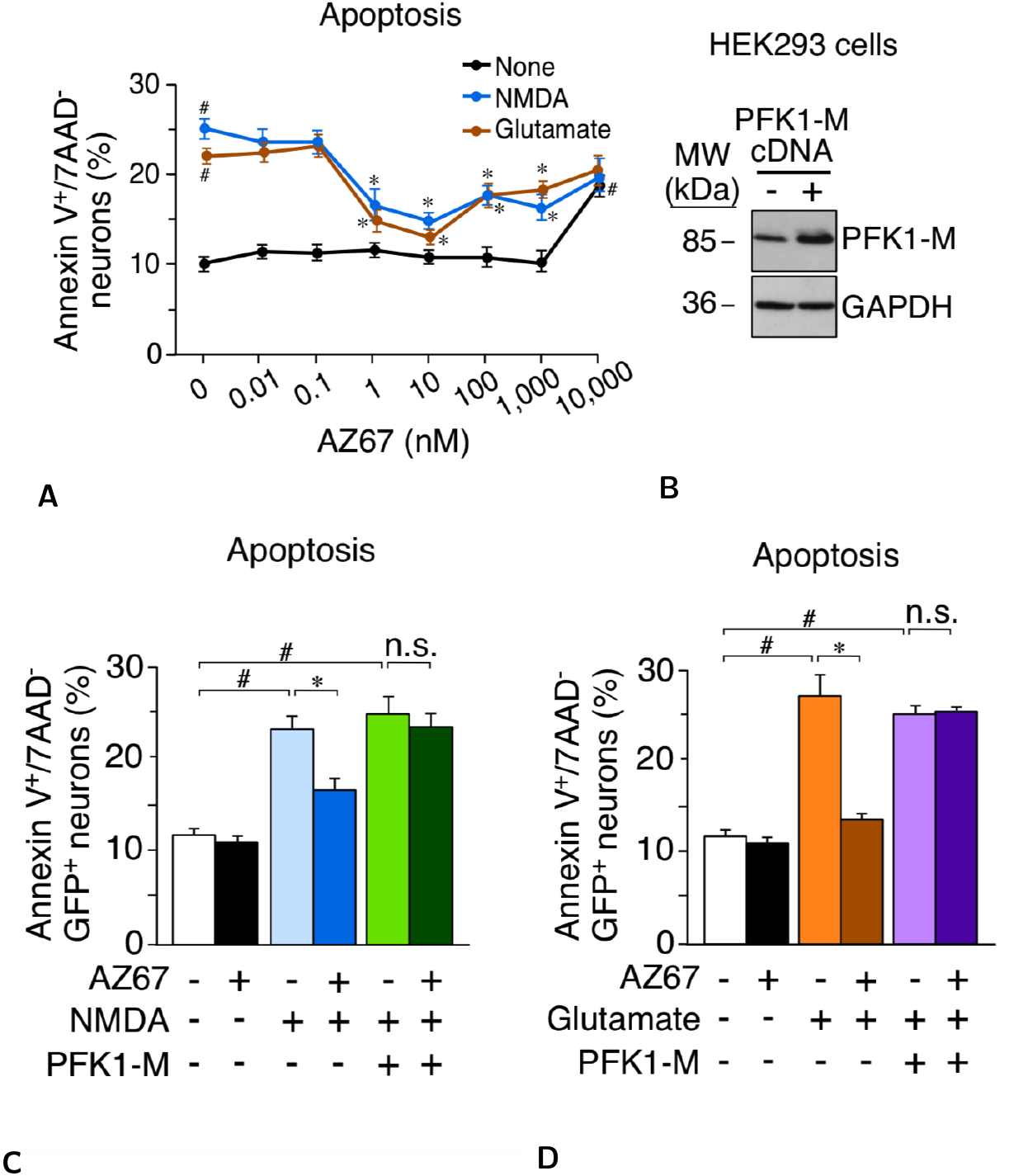
The neuroprotective effect of AZ67 is lost by abrogating glycolytic activation. **A** Treatment of neurons with NMDA (100 *µ*M; 10 mins) or glutamate (100 *µ*M; 10 mins), followed by washout, triggered apoptotic death after 24 h of incubation (compare NMDA or glutamate versus none values at 0 nM AZ67). Incubation of neurons with AZ67 for 24 h, after NMDA or glutamate was removed, dose-dependently prevented apoptotic death up to 10 nM, AZ67; at concentrations of 100 nM to 10 *µ*M, AZ67 showed progressive loss of protection. **B** Transfection of HEK293 cells with a plasmid vector expressing the full-length cDNA coding for the muscle isoform of PFK1 (PFK1-M), efficiently increased PFK1-M protein abundance, according to western blotting. **C** Treatment of neurons with NMDA (100 *µ*M; 10 mins), followed by washout, triggered apoptotic death after 24 h of incubation. Incubation of neurons with AZ67 (10 nM) for 24 h, after NMDA was removed, prevented apoptotic death. However, AZ67-mediated protection of apoptotic death was abolished when neurons were previously transfected with the full-length cDNA coding for PFK1-M. Apoptosis was analysed only in the efficiently-transfected, GFP+ neurons. **D** Treatment of neurons with glutamate (100 *µ*M; 10 mins), followed by washout, triggered apoptotic death after 24 h of incubation. Incubation of neurons with AZ67 (10 nM) for 24 h, after NMDA was removed, prevented apoptotic death. However, AZ67-mediated protection of apoptotic death was abolished when neurons were previously transfected with the full-length cDNA coding for PFK1-M. In all cases, data are mean ± S.E.M. values for n=3 independent culture preparations. #p*<*0.05 versus none at 0 nM AZ67; *p*<*0.05 versus the corresponding treatment at 0 nM AZ67 (ANOVA followed by the least significant difference multiple range test). n.s., not significant.

### *In vivo* AZ67 administration protects mice against the motor discoordination caused by a brain ischemia/reperfusion model

Finally, we aimed to investigate if AZ67 was able to exert neuroprotection in vivo. Since it is very well documented that brain injury in stroke occurs through an excitotoxic mechanism [29, 30], we studied whether AZ67 protected against damage caused in a mouse model of stroke. To achieve this, we induced a transient ischemia (30 min) by occlusion of the middle carotid artery (MCAO model), followed by 24 h reperfusion as described by a well-established protocol [31, 32]. AZ67 (60 mg/kg of body weight), or vehicle, were administered intravenously through the jugular vein immediately after the ischemic episode, at the start of the reperfusion. At this dose we found no toxicity 24 h after AZ67 administration in mice. Twenty-four hours after the transient MCAO episode, mice were subjected to the rotarod test, which revealed a ~ 40% performance of motor coordination when compared with the sham-operated animals (Figure 4A).

**FIG. 4:**
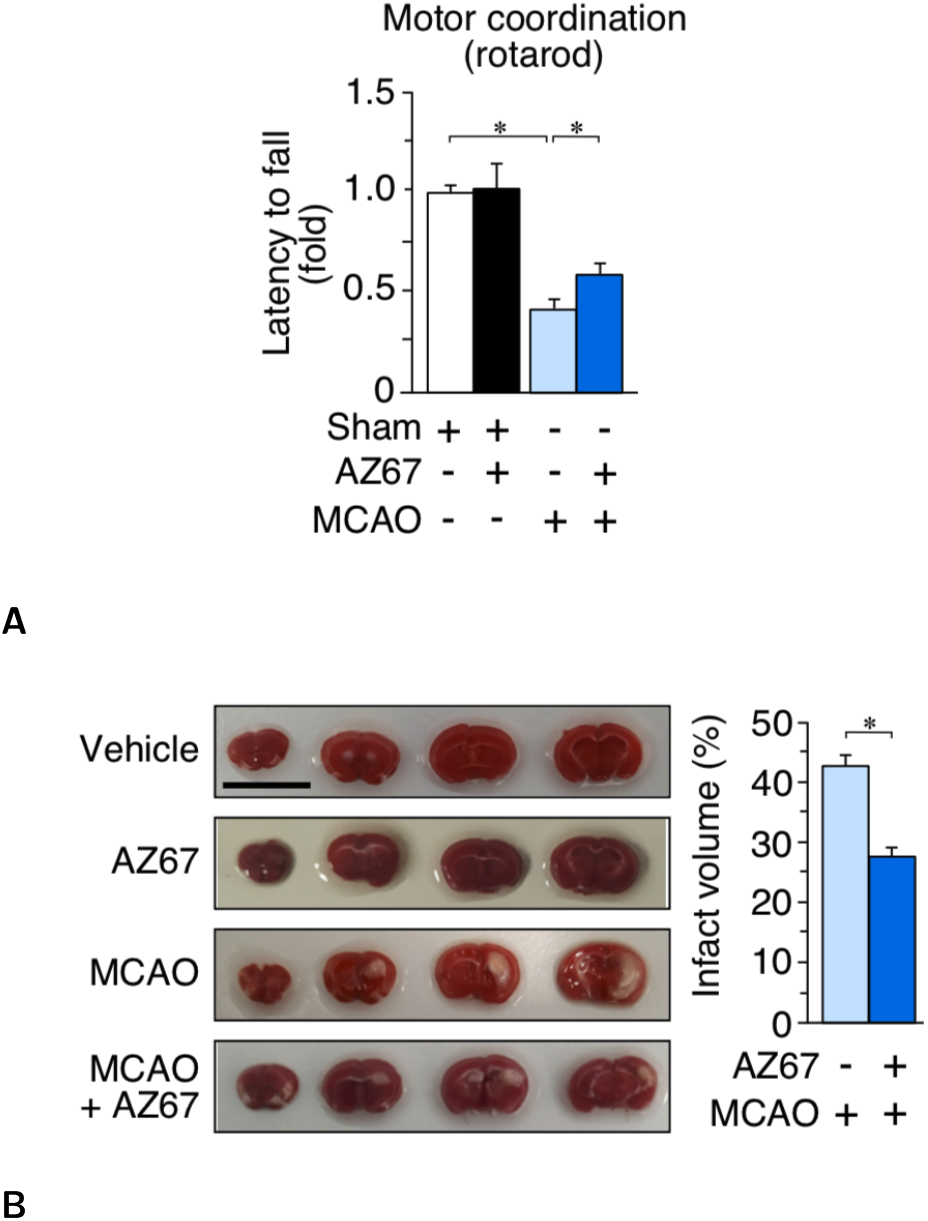
*In vivo* AZ67 administration protects mice against the motor discoordination and brain injury caused by a brain ischemia/reperfusion model. **A** Motor coordination, analysed 24 h after a transient MCAO episode in mice, revealed a ∼40% loss in performance (rotarod) when compared with the sham-operated animals. This effect was significantly prevented by the intravenous administration of AZ67 (60 mg/kg of body weight) immediately after the ischemic episode. **B** Infarcted brain volume, analysed 24 h after a transient MCAO episode in mice, was ∼40% of the brain of the sham-operated animals. This effect was significantly prevented by the intravenous administration of AZ67 (60 mg/kg of body weight) immediately after the ischemic episode. Left panel shows pictures of the brain sections of a representative animal for each experimental group. Bar, 1 cm. In all cases, data are mean S.E.M. values for 8 male mice. *p*<*0.05 (ANOVA followed by the least significant difference multiple range test).

Immediately after the test, mice were euthanized to determine the percentage of infarcted volume in the brain, which resulted to be ~43% (Figure 4B). Interestingly, administration of AZ67 significantly prevented the motor discoordination caused by MCAO, reaching a ~60% performance (Figure 4A), a result consistent with the significant protection of the infarcted brain volume, which decreased to 27% (Figure 4B).

## DISCUSSION

Here we show that pharmacological inhibition of PFKFB3 activity, by preventing glycolytic activation, exerts neuroprotection against excitotoxicity both in the NMDA-glutamate receptor activation model in primary cortical neurons and in the MCAO brain ischemic/reperfusion model *in vivo*. To our knowledge, this is the first time showing that inhibition of glycolysis, by the use of a small-molecule compound, shows a beneficial effect in a neurological disease model. In cancer cells, several PFKFB3 inhibitors of different chemical classes have been reported to inhibit glycolysis, on which these cells rely for proliferation and survival [33–37]. Amongst these PFKFB3 inhibitors, 3PO and its derivative PFK158 have been reported to reduce the cellular levels of F2,6BP, inhibit glucose uptake and lactate production, thus facilitating apoptosis in cancer cells. However, on our hands, PFK158 is inactive as PFKFB3 inhibitor in a purified human recombinant PFKFB3 enzymatic assay, at least at concentrations up to 100 *µ*M. Intriguingly, similar results were previously reported for 3PO [18, 37]. Whether these compounds inhibit glycolysis by interfering with a glycolytic target different to PFKFB3 remains to be elucidated. In contrast, AZ67 is a bona-fide PFKFB3 inhibitor [18] that, on our conditions, inhibited human recombinant PFKFB3 kinase activity at with an IC_50_ of 18 nM and decreased cellular F2,6BP production in A549 cells with an IC_50_ of 510 nM. When compared with astrocytes and other tissues, neurons are, by far, the type of cell showing the smallest PFKFB3 abundance, which is virtually absent [7]. In good agreement with this, we found that lower concentrations of AZ67 (in the 1-10 nM range) were sufficient to efficiently abrogate the enhancements in F2,6BP, glycolysis and apoptosis in neurons. Likewise, *in vivo* administration of the PFKFB3 inhibitor in the MCAO model could, in principle, enter all cells of the organism to broadly inhibit PFFKFB3 activity; however, it is highly likely that neuronal PFKFB3 would be the most sensitive against AZ67 given the particularly low PFKFB3 abundance in neurons [7], a feature that can be advantageous when determining the specific administration doses for future pre-clinical trials.

The rationale for targeting PFKFB3 and glycolysis as a therapeutic strategy for neurodegeneration relies on the physiological regulation of PFKFB3 protein stability in neurons. Thus, PFKFB3 is continuously degraded by the proteasome to keep glycolysis low in healthy neurons; however, PFKFB3 becomes stabilized upon proteasomal inhibition, leading to increased glycolysis and decreased PPP, which ultimately causes redox stress and neuronal death [7]. In good agreement with this notion, our data show that PFKFB3 inhibition by AZ67 exerts neuroprotection after blocking the proteasome with MG132. Furthermore, AZ67 prevented the neuronal death triggered by the Alzheimer’s disease-related peptide *Aβ*_25−35_, known to promote the degradation of Cdh1, i.e. the APC/C-cofactor necessary for PFKFB3 ubiquitylation and proteasomal degradation [22]. In the neuronal model of excitotoxicity, the increase in lactate release-resembling increased glycolysis-was paralleled with NADPH oxidation, which is a hallmark of decreased PPP. NADPH oxidation impairs glutathione regeneration thus causing redox stresss [7, 24], a feature that we have herein confirmed according to the increase mitochondrial ROS upon the excitotoxic insult. Interestingly, AZ67 was able to rescue the enhancement in lactate release, NADPH oxidation and redox stress, supporting the notion that restoring the equilibrium between glycolysis and PPP is a suitable and efficient neuroprotective strategy. Importantly, the neuroprotection exerted by AZ67 was abolished by over-expressing PFK1-M, an enzyme that activates glycolysis independently on F2,6BP levels [27]. This result demonstrates that the neuroprotective effect of AZ67 is due to its ability to inhibit the increase in glycolysis.

Excitotoxicity is a hallmark of various neurodegenerative diseases including stroke. Moreover, there is both pre-clinical [38] and clinical [39] evidence that tissue plasminogen activator (tPA), the only approved drug treatment for acute ischemic stroke, despite its obvious benefits presents adverse effects in the brain, including excitotoxicity potentiation [40]. Although neuroprotective strategies have been focused on NMDA receptor antagonists, unfortunately none of them have been successful, likely because functional NMDA receptors are essential for the normal brain physiology. Our data showing that targeting downstream the NMDA receptors activation - such as neuronal glycolysis-, rather than the NMDA receptors themselves, shows neuroprotection strongly suggest that targeting PFKFB3 would be a suitable alternative strategy for the prevention of the deleterious effects of NMDA receptor overstimulation in stroke and related excitotoxicity-associated neurological disorders, such as, amongst others, traumatic brain injury, Alzheimer’ or Parkinson’ diseases.

## METHODS

### PFKFB3 Enzymatic assay

Recombinant full length human PFKFB3 protein purified from Sf9 baculoviral system acquired from SignalChem (Cat. #P323-30G). ATP, fructose-6-phosphate and other chemicals were from Sigma-Aldrich. ADP detection system (ADP-Glo) was purchased from Promega. Inhibitors were synthesized as described by Boyd et al [18]. The kinase activity of the PFKFB3 protein was detected by measuring production of ADP from ATP in the presence of fructose-6-phosphate. The reactions were assembled in 384 well plates in a total volume of 25 *µ*l. Test compounds were serially diluted in dimethyl sulfoxide (DMSO). Reactions were set up by mixing test compounds with the enzyme and pre-incubating for 15 min. ATP and fructose-6-phosphate were next added to initiate the reactions. The final assay composition included: 100 mM Tris-HCl pH 8.0, 4 mM MgCl_2_, 5 mM KH_2_PO_4_, 5 mM DTT, 20 mM KF, 0.02% BSA, 1% DMSO (from the compounds), 15 nM enzyme, 20 *µ*M ATP (Km = 16 *µ*M) and 10 *µ*M fructose-6-phosphate (Km = 6 *µ*M). The kinase reactions were allowed to proceed for 1 hour at room temperature. Aliquots of the reaction mixtures (5 *µ*l) were transferred to fresh white 384 well plates and mixed with 5 *µ*l of the ADP-Glo reagent, followed by incubation for 30 min. The luminescent kinase detection reagent was added (10 *µ*l) and, following additional incubation for 15 min, the plates were read with a luminescence plate reader (Analyst HT). Positive (100%-inhibition) and negative (0%-inhibition) control samples were assembled in each assay plate and were used to calculate percent inhibition values of test compounds.

### Fructose-2,6-bisphosphate determinations

For F2,6BP determinations, cells were lysed in 0.1 N NaOH and centrifuged (20,000 × g, 20 min). An aliquot of the homogenate was used for protein determination, and the remaining sample was heated at 80°C (5 min), centrifuged (20.000 × g, 20 min) and the resulting supernatant used for the determination of F2,6BP concentrations using the coupled enzymatic reaction and F2,6BP standards as described by Van Schaftingen [41].

### Cell culture

Primary cultures of C57BL/6 mice cortical neurons were prepared from foetal animals of 14.5 days of gestation, seeded at 1.8 × 10^5^ cells/cm^2^ in plastic plates coated with poly-D-lysine (10 mg/ml) and incubated in Neurobasal (Life Technologies) supplemented with 2 mM glutamine, 5 mM of glucose, 0.25 mM pyruvate and 2% B27 supplement (Life Technologies). Cells were incubated at 37°C in a humidified 5% CO_2_-containing atmosphere. At 72 hours after plating, medium was replaced using Neurobasal (Life Technologies) supplemented with 2 mM glutamine, 5 mM glucose, 0.25 mM pyruvate and 2% B27 supplement (Life Technologies) minus antioxidants (MAO; i.e., lacking vitamin E, vitamin E acetate, superoxide dismutase, catalase and glutathione). Six days after plating medium was replaced again. Cells were used at day 9. Human embryonic kidney (HEK) 293 cells and adenocarcinomic human alveolar basal epithelial cells (A549 cells) were seeded at 10^4^ cells/cm^2^ in Dulbecco’s modified Earls Medium (DMEM; Sigma-Aldrich) supplemented with 10% fetal calf serum (FCS).

### Cell transfections

Cells (primary neurons or HEK293 cells) were transfected with 1.6 *µ*g/mL of a pIRES2-EGFP plasmid vector (Invitrogen) harbouring the full-length cDNA coding for the human muscle 6-phosphofructo-1-kinase muscle isoform (PFK1-M)^3^ (accession number, NM_000289.1) using Lipofectamine LTX-PLUS Reagent (Life Technologies) according with manufacturer’s protocol. Transfections were performed 24 hours before cells collection. Control cells were transfected with the empty vector.

### Western blotting

HEK293 cells were lysed in RIPA buffer (1% SDS; 2 mM EDTA; 12.5 mM Na_2_HPO_4_; 1% triton X-100; 150 mM NaCl; pH 7), supplemented with phosphatase (1 mM Na_3_VO_4_, 50 mM NaF) and protease (100 *µ*M phenylmethylsulfonyl fluoride (PMSF), 50 *µ*g/ml aprotinine, 50 *µ*g/ml leupeptine, 50 *µ*g/ml pepstatin, 50 *µ*g/ml anti-papain, 50 *µ*g/ml amastatin, 50 *µ*g/ml bestatin and 50 *µ*g/ml soybean trypsin inhibitor) inhibitor cocktail, 100 *µ*M phenylmethylsulfonyl fluoride and phosphatase inhibitors (1 mM o-vanadate). Samples were boiled for 5 min. Aliquots of cell lysates (50 *µ*g protein, unless otherwise stated) were subjected to sodium docedyl sulfate-polyacrylamide (SDS-PAGE) electrophoresis on an 8% acrylamide gel (MiniProtean, Bio-Rad) including PageRuler Plus Prestained Protein Ladder (Thermo). The resolved proteins were transferred electrophoretically into nitrocellulose membranes (Amersham protran premium 0.45 nitrocellulose, Amersham). Membranes were blocked with 5% (wt/vol) low-fat milk in 20 mM Tris, 150 mM NaCl, and 0.1% (w/v) Tween 20, pH 7.5, for 1 h. Subsequently, membranes were immunoblotted with anti-PFK1-M^4^ or anti-anti-GAPDH (4300, Ambion) primary antibodies overnight at 4°C. After incubation with horseradish peroxidase-conjugated goat anti-rabbit IgG (Santa Cruz Biotechnologies) (1/10000 dilution), membranes were incubated with the enhanced chemiluminescence kit WesternBright ECL (Advansta) before exposure to Fuji Medical X-Ray film (Fujifilm), and the autoradiograms scanned.

### Cell treatments

For NMDA receptors activation, neurons at 8 days in vitro were incubated with 100 *µ*M glutamate (plus 10 *µ*M glycine) or 100 *µ*M NMDA (plus 10 *µ*M glycine) for 10 minutes. Neurons were then washed and further incubated in culture medium with the PFKFB3 inhibitors for 24 hours. For amyloid-*β* treatment, the active truncated amyloid-*β* peptide *Aβ*_25−35_ (BioNova Cientifica S.L., Madrid, Spain) was used. *Aβ*_25−35_ was dissolved in distilled water at a concentration of 1 mg/ml and then incubated at 37°C for 3 days to induce its oligomerization [42]. We have shown that *Aβ*_25−35_ exerts neurotoxicity following identical mechanism to the full-length *Aβ*_1−42_ peptide [21]. Neurons were incubated in culture medium containing oligomerized *Aβ*_25−35_ (10 *µ*M) or the corresponding scramble non-aggregable peptide (*Aβ*_35−25_) (BioNova Cientifica S.L.), which was used as control. Neurons at 8 days *in vitro* were incubated with *Aβ*_25−35_ plus the PFKFB3 inhibitors for 24 hours. To inhibit the proteasome, neurons were incubated with MG132 (10 *µ*M) for 2 hours.

### Mitochondrial ROS

Mitochondrial ROS was detected using the fluorescent probe MitoSox™(Life Technologies). Cells were incubated with 2 *µ*M of MitoSox™for 30 minutes at 37°C in a 5% CO_2_ atmosphere in Hank’s Balanced Salt Solution (HBSS buffer); (NaCl 134.2 mM; KCl 5.26 mM; KH_2_PO_4_ 0.43 mM; NaHCO_3_ 4.09 mM; Na_2_HPO_4_ 2H_2_O 0.33 mM; glucose 5.44 mM; HEPES 20 mM; CaCl_2_ 2H_2_O 4 mM; pH 7.4). Cells were then washed with PBS (phosphate-buffered saline, 0.1 M) and collected by smooth trypsinization. MitoSox™fluorescence was assessed by flow cytometry and expressed in arbitrary units.

### Flow cytometric analysis of apoptotic cell death

APC/C-conjugated annexin-V and 7-amino-actinomycin D (7-AAD) (Becton Dickinson Biosciences, BDB, San Jose, CA, USA) were used to determine quantitatively the percentage of apoptotic neurons by flow cytometry. Cells were stained with annexin V-APC and 7-AAD, following the manufacturer’s instructions, and were analysed on a FACScalibur™flow cytometer (15 mW argon ion laser tuned at 488 nm; CellQuest software, Becton Dickinson Biosciences) using the CellQuest software (BDB). Both GFP^+^ and GFP^−^ cells were analyzed separately, and the annexin V-APC-stained cells that were 7-AAD-negative were considered to be apoptotic.

### Active caspase-3 determination

A fluorimetric caspase 3 assay kit (Sigma-Aldrich) was used following the manufacture’s protocol. This assay is based on the hydrolysis of the peptide substrate Ac-DEVD-AMC (acetyl-Asp-Glu-Val-Asp-7-amino-4-methylcoumarin) by caspase-3, which results in the release of fluorescent 7-amino-4-methylcoumarin (AMC). In brief, cells were lysed with 50 mM HEPES, 5 mM CHAPS, 5 mM DTT, pH 7.4 for 20 min on ice, and the assay buffer containing the Ac-DEVD-AMC substrate (20 mM HEPES, 2 mM EDTA, 0.1% CHAPS, 5 mM DTT, 16 *µ*M Ac-DEVD-AMC, pH 7.4) was added. Aliquots of 200 *µ*l were transferred to a 96-wells plate and the fluorescence recorded for 2 hours at 20 mins intervals at 37°C (*λ*exc=360 nm, *λ*em=460 nm). CSP-3 activity was determined as AMC release rate extrapolating the slopes to those obtained from a AMC standard curve. Results are expressed as fold change, arbitrarily assigning the value of 1 to control cells.

### NADPH/NADP**+** ratio determination

This was performed using the colorimetric NADPH/NADP assay kit (Abcam). Cells were resuspended in 500 *µ*l of NADPH/NADPH extraction buffer, vortexed and centrifuged at 14,000 rpm for 5 minutes to remove insoluble material. The supernatant was used for NADPH plus NADP measurement. NADPH was determined in 200 *µ*l of the supernatant, after heated at 60°C for 30 minutes to decompose NADP. Actual NADP and NADPH concentrations were calculated by extrapolating values to a NADPH standard curve (0-100 pmol/well).

### Lactate determination

Lactate released to the culture medium was determined as an estimation of glycolysis. To do so, the increments in absorbance of the culture medium samples were measured at 340 nm in a mixture containing 1 mM NAD^+^ and 22.5 units·ml^−1^ of lactate dehydrogenase in 0.25 M glycine/0.5 M hydrazine/1 mM EDTA at pH 9.5.

### Transient middle cerebral artery occlusion (MCAO)

Surgical endovascular insertion of a silicon-coated monofilament (602012PK10; Doccol Corporation, Sharon, MA, USA) was performed to induce transient middle cerebral artery occlusion (MCAO) for 45 minutes of ischemia, followed by filament removal to allow reperfusion [31, 32]. Briefly, 10-weeks-old C57BL/6J mice were anesthetized with sevoflurane (4% for induction, 3% for maintenance) in a mixture of O_2_/N_2_O (30/70%). After surgical exposure of the right carotid artery tree, the filament was inserted through the external carotid artery and advanced through the internal carotid artery until it reached the middle cerebral artery. The regional cerebral blood flow was monitored during surgery with a laser Doppler probe (Moor Instruments, Devon, UK). After 30 minutes of ischemia, the filament was removed to allow reperfusion. AZ67 (60 mg/kg of body weight) or vehicle were administered in a bolus (200 *µ*l) via the jugular vein immediately after reperfusion. Body temperature was maintained at 37 ± 0.5°C using a heating pad connected to a rectal probe (BAT-12 thermometer; Physitemp Instruments Inc., Clifton, NJ, USA). Mice were then sutured and returned to the cages. Sham-operated mice underwent the same surgical procedure without middle cerebral artery occlusion.

### Rotarod analysis

An accelerating rotarod test was used to determine motor coordination. Animals were trained during the immediate three previous days of the MCAO surgery. The first day, mice stayed on the rotating rod at a constant speed of 4 rpm, and the remaining 2^nd^ and 3^rd^ day they stayed at an accelerating speed (4 to 40 rpm in 5 mins). For the test, which was performed 24 hours after the MCAO surgery, mice were subjected to three consecutive trials at the accelerating speed for 5 mins (at 15 mins intervals). The latency to fall was determined and expressed in seconds.

### Infarct volume

Immediately after the rota-rod test, mice were euthanized by cervical dislocation after CO2 overdose, and the brain extracted and sliced in 2-mm coronal sections with a brain matrix on ice, which were used to determine the infarct volume after incubation of the slices in 2% (wt/vol) 2,3,5-triphenyltetrazolium chloride in phosphate-buffered saline (136 mM NaCl, 27 mM KCl, 7.8 mM Na_2_HPO_4_, 1.7 mM KH_2_PO_4_, pH 7.4) for 20 minutes at room temperature. Pictures of the brain sections were taken, and the images processed using the using the NIH image-processing package ImageJ 1.43n. Infarct volumes were determined by multiplying the selected infarcted area by the width of the slices. In order to correct the infarct volume by the edema, the ratio lesion volume of the ipsilateral (affected) versus that of the contralateral (unaffected) hemispheres was calculated. The percentage of infarct volume was calculated using the following formula: (infarcted volume corrected by edema × 100)/ Infarcted hemisphere volume.

### Statistical analysis

Results from cultured cells were obtained from 3 independent culture preparations using 4-6 technical replicates per sample. Data were expressed as mean ± standard error of the mean (SEM) values, using as ‘n’ the number of independent culture preparations. Statistical analysis of the results was performed by one-way analysis of variance (ANOVA), followed by the least significant difference multiple range test. In all cases, p*<*0.05 was considered significant. Statistics were performed using Microsoft Excel or the IBM SPSS Statistics software.

## ACKNOWLEDGEMENTS

JPB is funded by the MINECO (SAF2016-78114-R), CIBERFES (CB16/10/00282), H2020 European Commission (BatCure grant 666918) and FundaciÃşn Banco Bilbao-Vizcaya (FBBVA). AA is funded by H2020 European Commission (PANA grant 686009), Instituto de Salud Carlos III (PI15/00473; RD16/0019/0018 to AA) and Fundación Ramón Areces. JPB and AA are in part funded by FEDER. We acknowledge the technical assistance of Monica Resch, Monica Carabias-Carrasco, Lucia Martin and Estefania Prieto-Garcia from the University of Salamanca. OB, TE, DS, KZ and POF were funded by Gero Discovery LLC and acknowledge extensive technical assistance from and insightful discussions with Tim Pyrkov and Geny Getmantsev from Gero team. We appreciate S. Romanov and Nanosyn team for help with the synthesis and biological evaluation of PFKFB3 inhibitors.

## References

[1] J. P. Bolaños, A. Almeida, and S. Moncada, Trends in biochemical sciences 35, 145 (2010).

[2] J. Bolanos, S. Peuchen, S. Heales, J. Land, and J. Clark, Journal of neurochemistry 63, 910 (1994).

[3] A. Almeida, J. Almeida, J. P. Bolaños, and S. Moncada, Proceedings of the National Academy of Sciences 98, 15294 (2001).

[4] A. Almeida, S. Moncada, and J. P. Bolaños, Nature cell biology 6, 45 (2004).

[5] M. Delgado-Esteban, A. Almeida, and J. P. Bolaños, Journal of neurochemistry 75, 1618 (2000).

[6] P. GarciÌĄa-Nogales, A. Almeida, and J. P. Bolaños, Journal of Biological Chemistry 278, 864 (2003).

[7] A. Herrero-Mendez, A. Almeida, E. Fernández, C. Maestre, S. Moncada, and J. P. Bolaños, Nature cell biology 11, 747 (2009).

[8] P. Rodriguez-Rodriguez, E. Fernandez, and J. P. Bolaños, Journal of Cerebral Blood Flow & Metabolism 33, 1843 (2013).

[9] L. Pellerin and P. J. Magistretti, Journal of Cerebral Blood Flow & Metabolism 32, 1152 (2012).

[10] M. R. El-Maghrabi, F. Noto, N. Wu, and N. Manes, Current Opinion in Clinical Nutrition & Metabolic Care 4, 411 (2001).

[11] M. H. Rider, L. Bertrand, D. Vertommen, P. A. Michels, G. G. Rousseau, and H. Louis, Biochemical Journal 381, 561 (2004).

[12] C. Maestre, M. Delgado-Esteban, J. C. Gomez-Sanchez, J. P. Bolaños, and A. Almeida, The EMBO journal 27, 2736 (2008).

[13] P. Rodriguez-Rodriguez, E. Fernández, A. Almeida, and J. P. Bolaños, Cell death and differentiation 19, 1582 (2012).

[14] D. Lloyd-Jones, Circulation 119, 480 (2009).

[15] M. A. Moro, A. Almeida, J. P. Bolaños, and I. Lizasoain, Free Radical Biology and Medicine 39, 1291 (2005).

[16] J. Fan, T. M. Dawson, and V. L. Dawson, in Neurodegenerative Diseases (Springer, 2017) pp. 403–425.

[17] E. Sekerdag, I. Solaroglu, and Y. Gursoy-Ozdemir, Current neuropharmacology 16, 1396 (2018).

[18] S. Boyd, J. L. Brookfield, S. E. Critchlow, I. A. Cumming, N. J. Curtis, J. Debreczeni, S. L. Degorce, C. Donald, N. J. Evans, S. Groombridge, et al., Journal of medicinal chemistry 58, 3611 (2015).

[19] A. C. Klarer, J. O’Neal, Y. Imbert-Fernandez, A. Clem, S. R. Ellis, J. Clark, B. Clem, J. Chesney, and S. Telang, Cancer & metabolism 2, 2 (2014).

[20] S. Mondal, D. Roy, S. Sarkar Bhattacharya, L. Jin, D. Jung, S. Zhang, E. Kalogera, J. Staub, Y. Wang, W. Xuyang, et al., International journal of cancer 144, 178 (2019).

[21] R. Lapresa, J. Agulla, I. Sánchez-Morán, R. Zamarreño, E. Prieto, J. P. Bolaños, and A. Almeida, Neuropharmacology 146, 19 (2019).

[22] T. Fuchsberger, S. Martínez-Bellver, E. Giraldo, V. Teruel-Martí, A. Lloret, and J. Viña, Scientific reports 6, 31158 (2016).

[23] A. Almeida and J. P. Bolaños, Journal of neurochemistry 77, 676 (2001).

[24] O. Ben-Yoseph, P. A. Boxer, and B. D. Ross, Journal of neurochemistry 66, 2329 (1996).

[25] A. B. Knott, G. Perkins, R. Schwarzenbacher, and E. Bossy-Wetzel, Nature Reviews Neuroscience 9, 505 (2008).

[26] D. Nguyen, M. Alavi, K. Kim, T. Kang, R. Scott, Y. Noh, J. Lindsey, B. Wissinger, M. Ellisman, R. Weinreb, et al., Cell death & disease 2, e240 (2011).

[27] S. Vora, R. Oskam, and G. Staal, Biochemical Journal 229, 333 (1985).

[28] A. Almeida, J. P. Bolaños, and S. Moncada, Proceedings of the National Academy of Sciences 107, 738 (2010).

[29] J. Mallolas, O. Hurtado, M. Castellanos, M. Blanco, T. Sobrino, J. Serena, J. Vivancos, J. Castillo, I. Lizasoain, M. A. Moro, et al., Journal of Experimental Medicine 203, 711 (2006).

[30] Á. Chamorro, U. Dirnagl, X. Urra, and A. M. Planas, The Lancet Neurology 15, 869 (2016).

[31] O. Engel, S. Kolodziej, U. Dirnagl, and V. Prinz, Journal of visualized experiments: JoVE (2011).

[32] C. Rodríguez, M. E. Ramos-Araque, M. Domínguez-Martínez, T. Sobrino, I. Sánchez-Morán, J. Agulla, M. Delgado-Esteban, J. C. Gómez-Sánchez, J. P. Bolaños, J. Castillo, et al., Stroke 49, 2437 (2018).

[33] B. Clem, S. Telang, A. Clem, A. Yalcin, J. Meier, A. Simmons, M. A. Rasku, S. Arumugam, W. L. Dean, J. Eaton, et al., Molecular cancer therapeutics 7, 110 (2008).

[34] B. F. Clem, J. O’Neal, G. Tapolsky, A. L. Clem, Y. Imbert-Fernandez, D. A. Kerr, A. Klarer, R. Redman, D. M. Miller, J. O. Trent, et al., Molecular cancer therapeutics, molcanther (2013).

[35] T. V. Pyrkov, I. A. Sevostyanova, E. V. Schmalhausen, A. N. Shkoporov, A. A. Vinnik, V. I. Muronetz, F. F. Severin, and P. O. Fedichev, ChemMedChem 8, 1322 (2013).

[36] D. G. Brooke, E. M. van Dam, C. K. Watts, A. Khoury, M. A. Dziadek, H. Brooks, L.-J. K. Graham, J. U. Flanagan, and W. A. Denny, Bioorganic & medicinal chemistry 22, 1029 (2014).

[37] N. M. Gustafsson, K. Färnegårdh, N. Bonagas, A. H. Ninou, P. Groth, E. Wiita, M. Jönsson, K. Hallberg, J. Lehto, R. Pennisi, et al., Nature communications 9, 3872 (2018).

[38] Y. F. Wang, S. E. Tsirka, S. Strickland, P. E. Stieg, S. G. Soriano, and S. A. Lipton, Nature medicine 4, 228 (1998).

[39] R. Iglesias-Rey, M. Rodríguez-Yáñez, E. Rodríguez-Castro, J. M. Pumar, S. Arias, M. Santamaría, I. López-Dequidt, P. Hervella, C. Correa-Paz, T. Sobrino, et al., Translational stroke research 9, 347 (2018).

[40] R. Macrez, L. Bezin, B. Le Mauff, C. Ali, and D. Vivien, Stroke 41, 2950 (2010).

[41] E. Van Schaftingen, B. Lederer, R. Bartrons, and H.-G. Hers, European Journal of Biochemistry 129, 191 (1982).

[42] A. Almeida, J. P. Bolaños, and S. Moreno, Journal of Neuroscience 25, 8115 (2005).

